# The Mechanics of Temporal Interference Stimulation

**DOI:** 10.1101/2020.04.23.051870

**Authors:** Jiaming Cao, Brent Doiron, Chaitanya Goswami, Pulkit Grover

## Abstract

We utilize single neuron models to understand mechanisms behind Temporal Interference (TI) stimulation (also called “Interferential Stimulation”). We say that a neuron exhibits TI stimulation if it does not fire for a high-frequency sinusoidal input, but fires when the input is a low-frequency modulation of the high-frequency sinusoid (specifically that generated by addition of two high frequency sinusoids with a small difference in their frequencies), while the maximum amplitude is kept the same in both cases. Our key observation – that holds for both FitzHugh-Nagumo and Hodgkin-Huxley neuron models – is that for neuron models that do exhibit TI stimulation, a high frequency pure sinusoidal input results in a current balance between inward and outward currents. This current balance leads to a subthreshold periodic orbit that keeps the membrane potential from spiking for sinusoidal inputs. However, the balance is disturbed when the envelope of the sinusoids is modulated with a high slope: the fast-changing envelope activates fast depolarizing currents without giving slow outward currents time to respond. This imbalance causes the membrane potential to build up, causing the neuron to fire. This mechanistic understanding can help design current waveforms for neurons that exhibit TI stimulation, and also help classify which neuron-types may or may not exhibit TI stimulation.

---

^1^Transcranial electrical stimulation (TES) is widely used in neuroscientific studies and clinical treatments [1], [2], [3]. Several studies have examined techniques for improving resolution of TES, including using techniques from optimization. In [4], Grossman et al. used “Temporal Interference” (TI) stimulation (earlier called “Interferential Stimulation”, and applied largely to peripheral nervous system) which generated substantial interest (e.g. [5], [6], [7], [8]) due to its ability to stimulate deep inside the brain without shallow stimulation. This technique has been used earlier for transcutaneous stimulation (e.g. [9], [10]). The neural behavior that enables this is striking: a high-frequency sinusoidal input does not elicit firing, but a summation of two high frequency sinusoidal inputs with small differences in their frequencies fires neurons at the beat frequency [4]. This is true even when the maximum amplitudes of the two signals (pure sinusoid and summation of sinusoids) are kept equal. The concept generalizes in interesting ways to more than two sinusoids [6].

The recent work of Grossman et al. [4] as well as related efforts [9], [10], [4], [11] do not provide mechanisms of TI stimulation. Consequently, important questions about TI stimulation are not well understood, such as which neurons exhibit TI stimulation (see, e.g. [7]), or how to optimize parameters of TI stimulation for improved localization [6], [11]. This work provides the first understanding of the mechanics of TI stimulation using single neuron models. Specifically, we examine how ion-channel dynamics differ in response to interfering currents and pure sinusoidal currents, resulting in TI stimulation. With this understanding, spatiotemporal waveforms could be designed to trigger improved neurostimulation [6].

The key to understanding mechanisms of TI stimulation can be crystallized into the following succinct question: *why, for the same maximum amplitude, modulated high-frequency sinusoids lead to firing, but not pure sinuosids*. After all, the modulated signal has a lower average power than the pure sinusoid. Our key insight here is that, in essence, TI stimulation requires a *current balance*: for pure sinusoidal inputs, the inward and outward currents balance each other perfectly over each period of the input sinusoid. This current balance prevents a depolarization in the membrane potential from one cycle to the next, in turn preventing a neuron from firing. Instead, the neuron is maintained in a subthreshold periodic orbit.

This phenomenon can be thought of as an “envelope accommodation”. Classically, the term *neural accommodation* [12] is used to describe the phenomenon when a slow change in the input current’s amplitude (e.g., an input current with a small slope) does not cause a stimulation (see also [13], [14]). In our problem, a slow change in amplitude of the *signal’s envelope* does not cause firing. There, as is the case here, the accommodation results from balance between inward and outward currents that prevents the neuron from firing: a slow change in the current allows counter-polarizing currents to catch up with the fast depolarizing currents. The difference here is that this accommodation is happening in response to an envelope-modulated sinusoid. This is why this high-frequency current balance is not maintained at each time instant. Instead, only after averaging inward and outward currents over each cycle of the sinusoid does one see the current balance.

In Hodgkin-Huxley type neural models [13], [15], the current-balance in response to pure sinusoidal inputs arises from the activation and inactivation of sodium channels, as well as of the activation of potassium channels. A high-frequency inward current is generated by opening and closing of fast sodium channels (in direct response to the pure sinusoidal stimulus). This current is balanced *in each cycle of the sinusoid* by outward currents from relatively steady potassium channels (with some contribution from fast-changing leakage current). As a result, in response to high frequency pure sinusoidal input, the neural membrane potential displays a *stable subthreshold periodic orbit*, which resists firing for small changes in the current’s envelope. However, when the envelope is modulated substantially in a short amount of time, it leads to firing. This is because the sodium channels (that allow inward currents) respond to the modulation quickly, receiving a sharp boost, while the potassium channels (that allow outward currents) are slow to move from the steady-state they were at. The combination of sodium channels opening rapidly, and potassium channels being unresponsive, is what causes the neuron to spike.

Just as not all neural models exhibit classical accommodation, all neuron models do not exhibit TI stimulation (e.g. when a model does not attain the current balance described here). In such cases, when the neuron fires for modulated high-frequency sinusoids, it also fires for the same maximum amplitude pure sinusoid. In [7], we observed that, indeed, some neuron models (specifically, Parvalbumin (PV)-expressing inhibitory cells [16]) do not exhibit TI stimulation, whereas models of Hodgkin-Huxley squid neurons and excitatory pyramidal cells do exhibit TI stimulation. We anticipate that there are biological neurons that do not exhibit TI stimulation, and this work might suggest neurons which are good candidates for exhibiting (or not exhibiting) TI stimulation.

We discuss below why simplistic explanations such as low-pass filtering, or even a more sophisticated (and nonlinear) envelope-demodulation reasoning, are insufficient to explain temporal interference.

## Insufficiency of a low-pass filtering or a linear resonant neuron explanation

A sub-threshold low-pass filtering or linear resonance [17] by itself cannot lead to TI stimulation. This is because any linear transform on a summation of two high-frequency sinusoids would yield the same result as a single sinusoid of high frequency (and twice the amplitude). After all, both signals reside in high frequencies, and ideal low pass filtering/resonant filtering either of them (and their sum) will yield 0. To utilize the *envelope modulation* of the sinusoid, a nonlinearity of the system needs to be exploited. This leads us to the second explanation, suggested in [4], [18].

## Insufficiency of simplistic amplitude-demodulation explanation

One appealing explanation is that neurons somehow perform an envelope demodulation, and fire when the demodulated signal exceeds a threshold. This is the explanation appealed to in [4], with minor modifications in [18]. For such envelope demodulation, one indeed needs nonlinearity. E.g., a simple envelope demodulation circuit has a diode followed by a low-pass filter made of a capacitor and resistor in parallel (see, e.g. [19]). Indeed, there is a diode-like behavior in neural dynamics, namely that of sodium currents, which tend to flow only in one direction because the sodium Nernst potential is high. Sodium channels, as we observe in Fig. 4, also respond to high frequencies, explaining why neurons respond at all to amplitude modulated sinusoidal inputs. Thus, one might think that temporal interference stimulation works by charging the neural membrane by affecting only sodium channels. While this is definitely a part of our reasoning, it is insufficient by itself. This is because this explanation would hold even when pure sinusoidal inputs are used, and thus it fails to explain why the experimentally examined neurons in [4] do not fire with a pure sinusoidal input. The key is to explain why pure sinusoidal stimuli do no stimulate, but a sum of two high frequency sinusoids can.

**Fig. 1.**
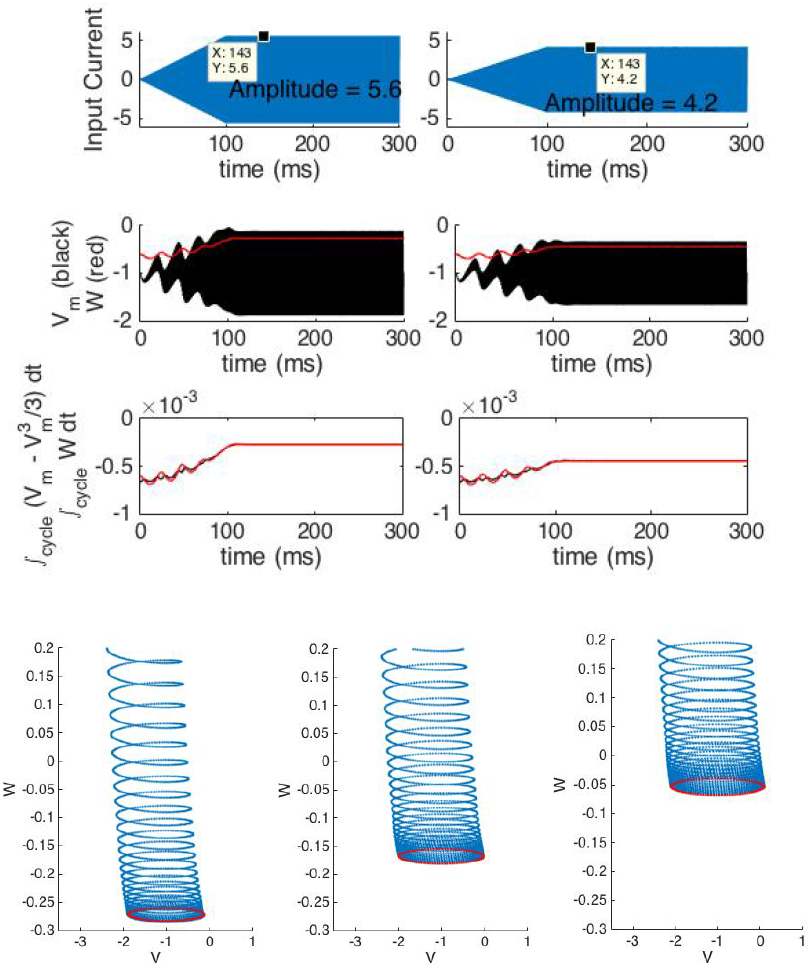
(Top figure; top row) Sinusoidal current inputs with amplitudes ramped up to reduce effects of initial state. (Top figure; middle row) Response to the sinusoidal current with ramped amplitude shows that no stimulation is observed. (Top figure; bottom row) Examining integrated *V_m_* and *W* currents over one cycle of the sinusoid. While an initial mismatch in currents leads to a firing event, the currents settle down and match each other exactly over a cycle, thereby canceling each other and keeping membrane potential’s DC-value a constant. A frequency of 1000 Hz is used for the base sinusoid.(a) Shows this balance for a larger and (b) for a smaller value of the current. (Bottom) For pure sinusoidal inputs, the (V, W)-phase space approaches a stable subthreshold periodic orbit (shown in red). The amplitude of the currents varies from 5.6 (left), 6.3 (middle) to 7 (right). At these periodic orbits, the currents are balanced in each cycle: the W-current balances the *V_m_*-current perfectly

**Fig. 2.**
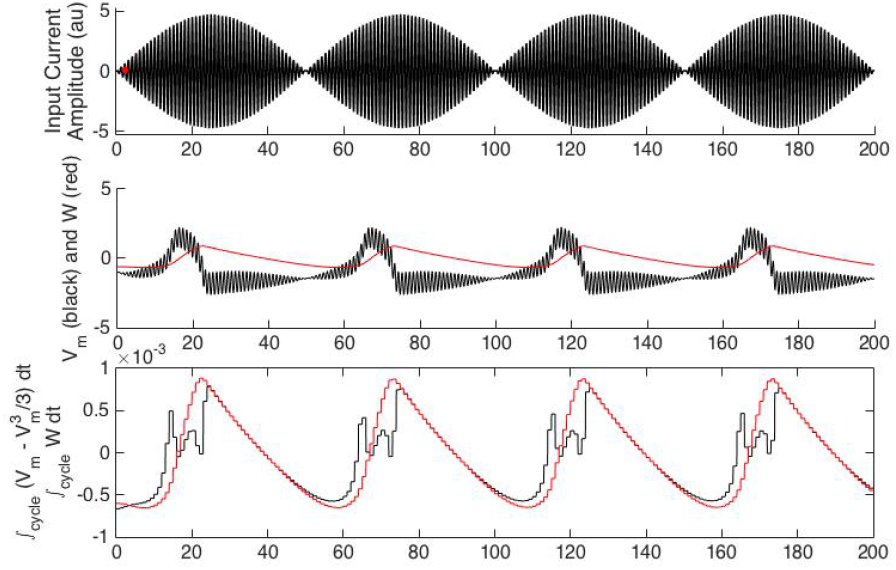
Amplitude modulated input currents can cause stimulation. The input current is a sum of two sinusoidal inputs of frequencies 1000 and 1010 Hz, and equal amplitude. The lower most plot examines the integrated *V_m_* and *W* currents over one cycle of the base sinusoid (1005 Hz). Current mismatches for extended periods of time lead to neural firing. In particular, the *V_m_*-current leads the W-current (which is slow to follow), causing the membrane potential to spike. Note that the total amplitude of currents is the same in Figs. 1 and 2.

**Fig. 3.**
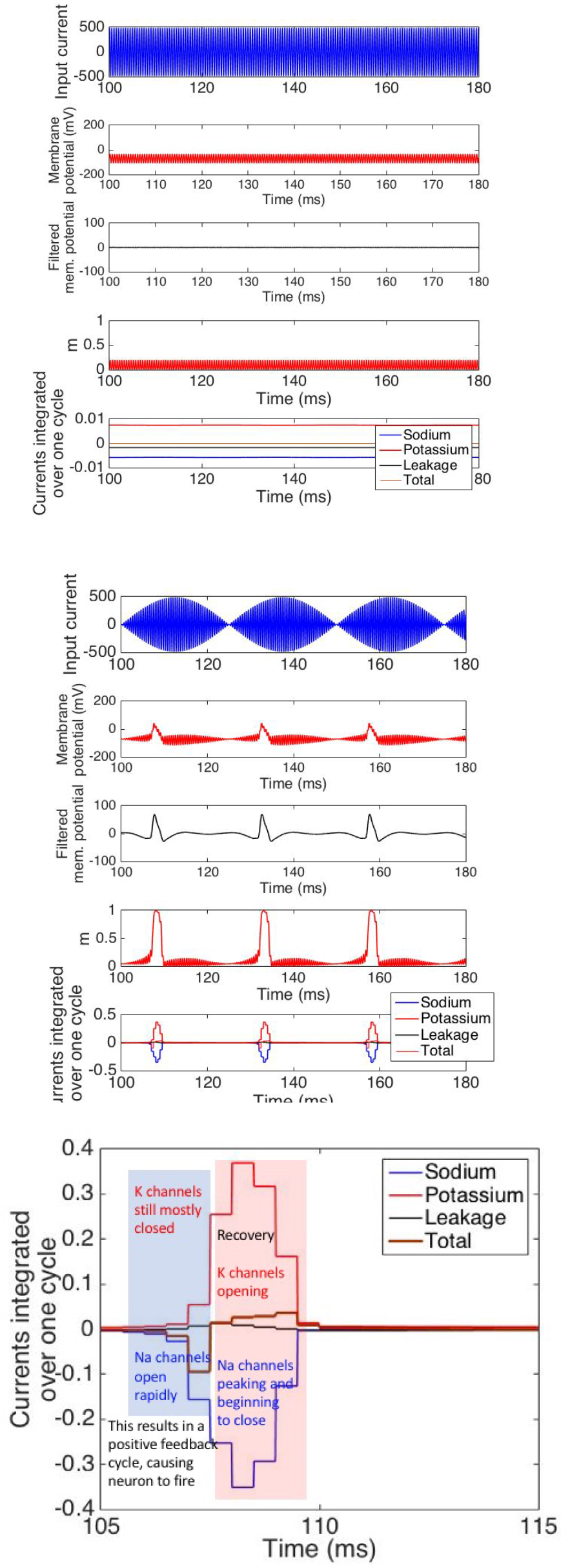
Illustration of current balance for pure sinusoidal inputs for neurons that exhibit Temporal Interference stimulation. The top figure, for pure sinusoidal input, shows no stimulation (*V_m_* stays near −70 mV, pass-band filtered membrane potential is zero, and sodium activation, *m*, stays small). This is because of the current balance, shown in the last subplot of the top figure. Integrated over a cycle, sodium, potassium, and leakage currents cancel each other exactly. In the middle figure, a modulated sinusoidal input stimulates the neuron into firing. Besides the *V_m_* and filtered Vm evidence, the most direct evidence is m rising to its peak value, 1. The current balance is disturbed with a strong and quick swing towards sodium current (negative in this figure) before potassium current catches up. A zoomed-in version of this short-time imbalance is shown in the lowermost figure.

**Fig. 4.**
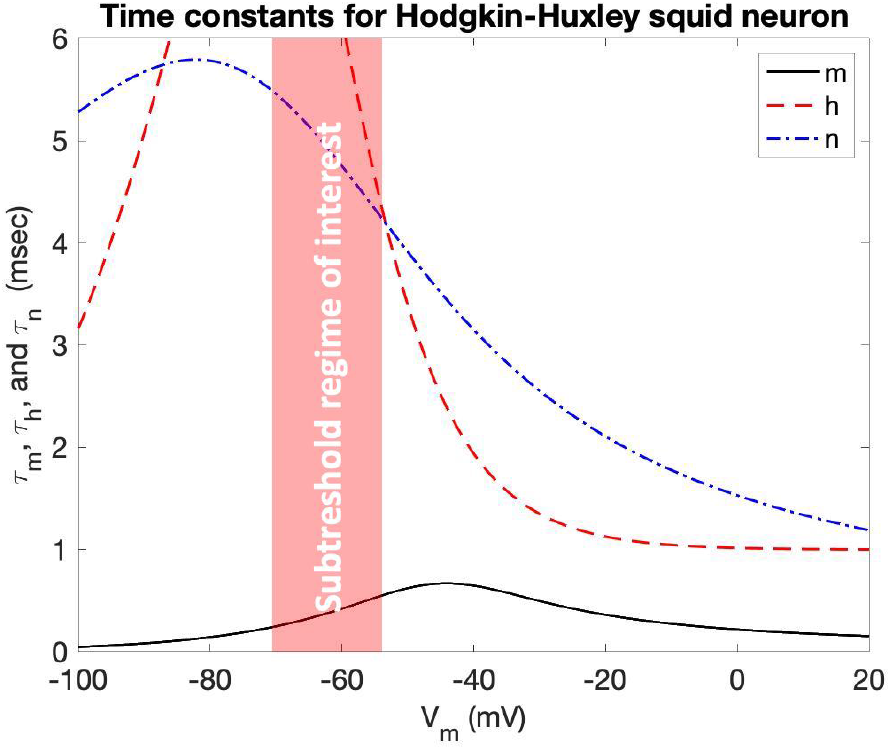
The *m, n, h* time-constants of the Hodgkin-Huxley neuron as they vary with *V_m_*. The regime of most interest is the subthreshold regime close to −65 mV, the resting membrane potential of the Hodgkin-Huxley neuron. In this regime, *τ_h_* and *τ_n_* are more than ~8x to 12x slower than *τ_m_*, creating the

Finally, our results have practical implications on optimizing waveform design for TI stimulation. E.g., in [18], the authors suggest that to maximize firing “envelope amplitudes”^2^ should be maximized. Our current-balance hypothesis suggests that the slope of the envelope plays a critical role, and maximizing amplitude of the envelope itself is not sufficient (e.g., the amplitude could be made large with a small slope by performing a very low frequency modulation of a large amplitude sinusoid).

## I. Models and dynamical systems equations

### Integrate-and-Fire models

The IF-type models that we examined are leaky integreate and fire (LIF) and quandratic/exponential integrate and fire (QIF and ExpIF respec-tively; see [15] for definitions).

### FitzHugh-Nagumo model

We use the FitzHugh-Nagumo model with dynamics as described by two parameters, *V_m_* and *W*, below:

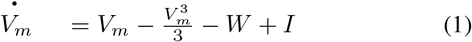

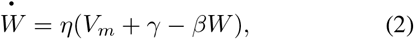

for some constants *η, β* and *γ*, with *η* typically being small to reflect slow dynamics of *W* relative to dynamics of *V_m_*. We observe TI stimulation for varying values of these parameters. The plots are obtained for *η* = 0.08, *γ* = 1, and *β* = 0.01. We refer to the term 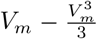 as “*V_m_*-current,” and *W* is called the *W*-current.

### Hodgkin-Huxley models

We use the classic 4-dimensional Hodgkin-Huxley (HH) model. Parameters are used for the original Hodgkin-Huxley squid neuron [13], although we have earlier observed that TI stimulation is observed in some other HH-type models as well [7]. General equations are described as follows:

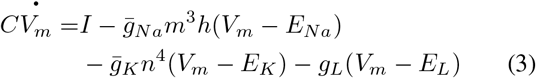

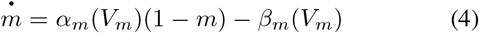

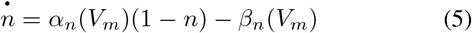

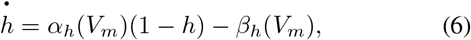

The · on top of a variable denotes the time derivative, *α*(*V_m_*)’s and *β*(*V_m_*)’s are functions of *V_m_* that do not change with time, and for the HH model are given by:

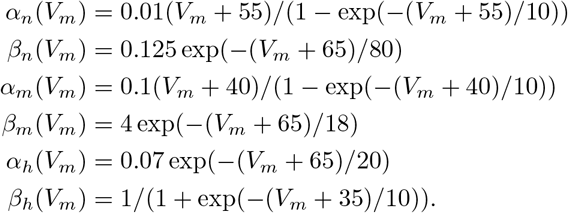

*m, n*, and *h* are gating variables that take values between 0 and 1, and correspond to sodium activation, potassium activation, and sodium deactivation gates respectively. The probability that the sodium activation gate is open is *m*^3^, the probability that the sodium inactivation gate is open is *h*, and the probability that a potassium channel is open is *n*^4^. 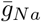 and 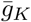 are maximum conductances for these channels, *E_Na_, E_K_*, and *E_L_* are the reversal potentials for each channel. The steady-state values of *m, n* and *h* variables, as functions of the membrane potential, are denoted by *m*_∞_, *n*_∞_ and *h*_∞_. These are given by: *m*_∞_ = *α_m_*/(*α_m_*+*β_m_*), *n*_∞_ = *α_n_*/(*α_n_* + *β_n_*), and *h*_∞_ = *α_h_*/(*α_h_* + *β_h_*). Also, the time constants that describe the dynamics of the *m, n*, and *h* parameters at any voltage *V_m_* are given by: *τ_m_*(*V*) = 1/(*α_m_* + *β_m_*), *τ_n_*(*V*) = 1/(*α_n_* + *β_n_*), and *τ_h_*(*V*) = 1/(*α_h_* + *β_h_*).

### Model of how intracellular current is generated by extracellular stimulation

Our understanding of mechanics of TI stimulation is built on the understanding of how extracellular currents induce changes in membrane potential that can cause neurostimulation. In this work, we use single compartment models, and assume that extracellular currents around the neuron induce proportional intracellular currents in the neuron. That is, a multiplicative factor connects extracellular and intracellular currents. This is justified by evidence in the literature, e.g., the work of Rattay [20] that shows that the induced current in an axon is proportional to the spatial derivative of the extracellular current parallel to the axon. Because the temporal structure of the waveform is preserved, for our simple single compartment model, the induced intracellular currents can be assumed to be proportional to the extracellular currents.

### Estimating firing events

A nontrivial issue is examining waveforms for estimating when a neuron fires. For FitzHugh-Nagumo and Hodgkin-Huxley models, since there is no fixed threshold that the membrane potential should exceed, we determine neuron firing by observing a sharp increase in the amplitude of the response of the membrane potential in response to a small increase in applied current waveform amplitude (a multiplicative factor for the entire waveform). Any response with amplitude beyond this sharp jump is considered to be neuron firing. For further examination, the shape of the spike is examined after filtering the signal to retain the spiking. Finally, for Hodgkin-Huxley neurons, a secondary validation for firing can be obtained by examining values of the parameter m. If m approaches 1, it reliably indicates firing.

## II. IF-type models do not exhibit TI stimulation

IF-type models, linear and nonlinear, do not exhibit TI stimulation because both sinusoidal and envelope-modulated inputs cause stimulation. The models that we tested include Leaky-Integrate and Fire (LIF), Quadratic Integrate and Fire (QIF), and Exponential Integreate and Fire (ExpIF). Despite varying parameter choices (frequencies and amplitudes of the sinusoids) over wide ranges, none of these models exhibit temporal interference stimulation. This is because in absence of a subthreshold polarizing current, there is no current to balance the depolarizing current for a pure sinusoidal input. All three models respond in qualitatively similar ways: at maximum amplitudes that cause firing, both modulated and unmodulated sinusoidal inputs cause the cells to fire. This is not surprising: as long as enough current is integrated over, the neuron fires. Once it fires, the neuron is set to the same initial state, leading to repeated firing.

## III. TI Stimulation and current balance in FitzHugh-Nagumo models

We observed that FitzHugh-Nagumo models exhibit TI stimulation only for a subset of possible parameter values. Thus, to understand the aspects responsible for TI stimulation, we deliberately chose FitzHugh-Nagumo model parameters that do exhibit TI stimulation. For these choices, pure sinusoidal inputs may cause an initial spiking of activity (depending on the neuron’s initial state; to sidestep this issue, in Fig. 1 we use a ramped up sinusoid, as is also done in [4]). Regardless, the neural state for a pure sinusoidal input converges to a stable subthreshold periodic orbit in the phase space. To confirm that the neuron has not fired, we examine the values of *V_m_* and W, both of which remain small (see Fig. 2 for comparison). This convergence to a periodic orbit for (V, W) parameters of a FitzHugh-Nagumo neuron is illustrated in Fig. 1 (bottom) for three different values of the input, where the orbit itself is also illustrated (in red). The time the state takes to circle once along this periodic orbit coincides with the period of the input sinusoid. That is, at the end of each cycle, both *V_m_* and W return to the same point. In turn, this is a consequence of the current balance – when integrated over one cycle of the sinusoid – between the (inward) *V_m_* and the (outward) W currents for this neuron. To confirm the existence of this average current balance, we plot the integrated the currents over each cycle in Fig. 1 (lower plot in each sub-plot). Both *V_m_* and W currents, integrated over a cycle, exactly coincide in the steady state, canceling each other out. Thus, the membrane potential does not rise in any given cycle of the sinusoid for a pure sinusoid input.

### Why sudden and large change in the envelope of a high frequency sinusoidal input current makes the neuron fire

We plot the parameters *V_m_* and *W* vs time in Fig. 2 for a sum of two high-frequency sinusoidal currents of equal amplitudes (1000 ad 1010 Hz). As is illustrated in Fig. 2 (bottom), when the envelope of the *V_m_* current rises, the current balance is disrupted for summation of two sinusoidal inputs because the dynamics of *V_m_*-current are faster than that of the *W*-current (and are affected by high frequencies). The current balance between (average) *V_m_*-current and *W*-current that was maintained for a pure sinusoidal input is no longer maintained because while *V_m_* current responds quickly to the increasing envelope, *W*-current responds, but only after some lag, and this lag creates an increase in *V_m_* that is sufficient to make the neuron fire. The firing is detected by a) observing that the membrane potential exceeds 0, and is substantially higher than that in Fig. 1; b) The shape of the spike is consistent with the spike shape for FitzHugh-Nagumo model (see, e.g. [15], [21]); c) There is a sharp increase in amplitude of the detected spike in response to increase in input current by a multiplicative factor (across time); and d) The parameter W also takes large values after the detected spiking event.

Sub-threshold oscillatory behavior of *V_m_* is resumed (temporarily) after the firing event, and this is again caused by a (temporary) current balance (e.g. between 30 and 50 ms on x-axis in Fig. 2). Once again, when the envelope of the input current starts rising, the neuron fires again (e.g. between 60 and 75 milliseconds). Thus, there is no stable subthreshold periodic orbit for this modulated-envelope input.

## IV. TI Stimulation and Current Balance in Hodgkin-Huxley models

In Hodgkin-Huxley models, we chose the classic squid giant axon [13] because of the textbook nature of the model, although other neuron models also exhibit TI stimulation (while some do not) [7]. For this model, TI stimulation is clearly observed above 1700 Hz. That is, a sum of two sinusoids with a small difference in their frequencies, e.g., 10 Hz, we observe stimulation, while a single sinusoid of 1700 Hz and twice the amplitude does not exhibit TI stimulation. For lower frequencies, especially lower than 1400 Hz, we did not observe TI stimulation, despite sweeping the parameter space. We now examine the mechanics of channels to understand what causes TI stimulation.

### Pure sinusoid inputs

For pure sinusoidal inputs, a similar current balance is observed for the Hodgkin-Huxley neuron (Fig 3) (top). The neuron does not fire for a high-frequency pure sinusoidal input, as evidenced by a) low membrane potential; b) small filtered membrane potential; c) small values of *m* parameter attained (*m* typically rises to 1 when the neuron fires). Indeed, a current balance is again implicated in the neuron not firing. For this neuron model, the balance is between 3 currents, namely, sodium, potassium, and leakage currents, all integrated over one cycle of the sinusoid. To illustrate that the currents are balanced, a ‘Total’ current, integrated over each cycle, is also shown, which is a sum of these three currents. This ‘Total’ current (integrated over each cycle) is precisely zero for pure sinusoidal inputs, which implies that the membrane potential does not change from one cycle to another.

### Envelope-modulated inputs

The response of various gating parameters to amplitude-modulated sinusoidal input currents is shown in Fig. 3 (middle; and zoomed-in version at the bottom). In essence, the FitzHugh-Nagumo reasoning extends: fast sodium current responds to the increasing envelope. Sodium channels, which are responsible for the depolarizing currents here, open to let in currents and increase the membrane potential. Changes in amplitude of the input sinusoid lead to larger and larger currents being driven into the cell in every cycle (see zoomed-in version in the lower most plot in Fig. 3). Potassium currents are unable to adapt quickly to these increases in membrane potential (owing to their large time constants), leading to further increase in membrane potential that quickly leads to the neuron getting into the positive feedback cycle that leads to firing. Potassium current catches up, but at that point the neuron is already in the positive feedback cycle leading to firing. This lag creates a short-time current imbalance that can be observed in Fig 3 (bottom), that depolarizes the membrane.

This current balance is observed for all models that exhibit TI stimulation. It is also worth noting that for neurons that do *not* exhibit TI stimulation, for high enough sinusoid amplitude, the current balance does not hold, and the potential accumulates over several cycles. Indeed, it is shown in [7] that this neural model does not exhibit TI stimulation.

### Why do ion-channels respond to these high-frequency currents?

Conventional wisdom suggests that these high-frequency currents, operating at 1000s of Hz, will not affect the neuron. However, as we see in both FitzHugh-Nagumo and HH models, even when a pure sinusoidal current is input, it is a very active cancellation that makes a neuron not fire. Both depolarizing and polarizing currents seem to respond to the high frequency stimulating currents.

We illustrate the mechanism behind this by focusing on the HH neuron. How can sodium channels respond to such high frequencies? This is because the time constants for sodium channels are generally quite low, as shown in Fig. 4. In fact, *there is a direct link between the sodium-channel activation time constant and the carrier frequency of the amplitude modulated signal that results in TI stimulation*. For all values of the membrane potential *V_m_*, in this model, this time constant, *τ_m_*, is smaller than 0.6 milliseconds. This corresponds to a center frequency of about 1700 Hz, which is on the lower end of the frequency range when the neuron starts exhibiting TI stimulation. Thus, sodium channel time constants can inform the carrier frequency in TI stimulation.

The high-frequency sinusoidal current opens and closes sodium channels. Sodium currents are (largely) one-directional, going into the cell, because *E_Na_*, the Nernst potential of sodium, is positive and large. This means that the high-frequency sinusoid charges the neural membrane, depolarizing it. This depolarization is still somewhat slow, especially at low input amplitudes, because the membrane capacitance is being charged by a rather high-frequency input.

Why do potassium channels respond at all? Indeed, potassium channels’ gating parameter, *n*, does not respond directly to the stimulating current because of its large time constant. However, the slow increase in membrane potential due to opening and closing of sodium channels causes the potassium channels to open and close as well because they are voltage gated (and thereby changing *n* as well).

## V. Varying neural parameters to observe changes in current parameters for TI stimulation

To test our explanation of the mechanics of TI stimulation, in this section, we vary neural parameters, and observe if the neuron’s exhibition of TI stimulation reflects predictions based on our explanation above. First, by slowing down the sodium activation parameter, *m*, we expect TI stimulation to occur at even lower frequencies. This is exactly what we observe. In Fig. **S1** (see Supplementary Information), we slowed down *m* by increasing its time constant so that the original time constant is 75% of the new time constant (across all values of *V_m_*). The resulting neuron exhibits TI stimulation at 1200 Hz, whereas the classical Hodgkin-Huxley neuron exhibits TI stimulation only above 1600 Hz.

By slowing down the sodium activation, we also expect changes in the required slope of the envelope to cause stimulation. Specifically, slower sodium channels would find it hard to respond to slower envelopes: the membrane potential would rise too slowly for sodium channels to drive the cell to excitation before potassium channels open. This is indeed observed in our computational model as well, as illustrated in Fig. **S2** in Supplementary Information. At sinusoidal frequencies of [1600,1633] Hz, and amplitudes of 187.27, while the classical Hodgkin-Huxley neuron fires, a version of the neuron that has sodium channels slowed down (as above) does not fire. Below this amplitude, and below this difference in frequencies (keeping the lower frequency as 1600 Hz), neither neuron fires (note that it is the small difference in frequencies that keeps the envelope slope small).

Similarly, speeding up sodium inactivation parameter, *h*, or potassium activation parameter, *n*, is expected to affect the

Qualitatively, these experiments demonstrate consistency of our predictions with the observations.

neuron so that it does not exhibit TI stimulation at this frequency and amplitude. Indeed, this is exactly what happens, as is shown in Fig. **S3** in Supplementary Information.

## Footnotes

1 Author order alphabetical. This material is based upon work supported by the Naval Information Warfare Center (NIWC) Atlantic and the Defense Advanced Research Projects Agency (DARPA) under Contract No. N65236-19-C-8017. Any opinions, findings and conclusions or recommendations expressed in this material are those of the author(s) and do not necessarily reflect the views of the NIWC Atlantic and DARPA.

2 This envelope amplitude is defined in [18] as the difference between the peak and the minimum value of the envelope.

## REFERENCES

[1] Michael A Nitsche and Walter Paulus. Excitability changes induced in the human motor cortex by weak transcranial direct current stimulation. The Journal of physiology, 527(3):633–639, 2000.

[2] Djamila Bennabi and Emmanuel Haffen. Transcranial direct current stimulation (tDCS): a promising treatment for major depressive disorder? Brain sciences, 8(5):81, 2018.

[3] Michael A Nitsche, Leonardo G Cohen, Eric M Wassermann, Alberto Priori, Nicolas Lang, Andrea Antal, Walter Paulus, Friedhelm Hummel, Paulo S Boggio, Felipe Fregni, et al. Transcranial direct current stimulation: state of the art 2008. Brain stimulation, 1(3):206–223, 2008.

[4] Nir Grossman, David Bono, Nina Dedic, Suhasa B Kodandaramaiah, Andrii Rudenko, Ho-Jun Suk, Antonino M Cassara, Esra Neufeld, Niels Kuster, Li-Huei Tsai, Alvaro Pascual-Leone, and Edward S. Boyden. Noninvasive deep brain stimulation via temporally interfering electric fields. Cell, 169(6):1029–1041, 2017.

[5] Sumientra Rampersad, Biel Roig-Solvas, Mathew Yarossi, Praveen P Kulkarni, Emiliano Santarnecchi, Alan D Dorval, and Dana H Brooks. Prospects for transcranial temporal interference stimulation in humans: a computational study. NeuroImage, 202:116–124, 2019.

[6] Jiaming Cao and Pulkit Grover. Stimulus: Noninvasive dynamic patterns of neurostimulation using spatio-temporal interference. IEEE Transactions on Biomedical Engineering, 2019.

[7] Jiaming Cao and Pulkit Grover. Do single neuron models exhibit temporal interference stimulation? In 2018 IEEE Biomedical Circuits and Systems Conference (BioCAS), pages 1–4. IEEE, 2018.

[8] Jacek Dmochowski and Marom Bikson. Noninvasive neuromodulation goes deep. Cell, 169(6):977–978, 2017.

[9] GC Goats. Interferential current therapy. British journal of sports medicine, 24(2):87, 1990.

[10] Brenda Savage, AG McC, and John R Roberts. Interferential therapy. Faber & Faber, 1984.

[11] Yu Huang, Abhishek Datta, and Lucas C Parra. Optimized interferential stimulation of human brains. bioRxiv, page 783423, 2019.

[12] Keith Lucas. On the rate of variation of the exciting current as a factor in electric excitation. The Journal of physiology, 36(4-5):253, 1907.

[13] Alan L Hodgkin and Andrew F Huxley. A quantitative description of membrane current and its application to conduction and excitation in nerve. The Journal of physiology, 117(4):500–544, 1952.

[14] AB Vallbo. Accommodation related to inactivation of the sodium permeability in single myelinated nerve fibres from xenopus laevis. Acta physiologica scandinavica, 61:429–444, 1964.

[15] Eugene M Izhikevich. Dynamical systems in neuroscience. MIT press, 2007.

[16] Balázs Chiovini, Gergely F Turi, Gergely Katona, Attila Kaszás, Dénes Palfi, Pal Maak, Gergely Szalay, Mátyás Forián Szabó, Gábor Szabó, Zoltán Szadai, et al. Dendritic spikes induce ripples in parvalbumin interneurons during hippocampal sharp waves. Neuron, 82(4):908–924, 2014.

[17] Bruce Hutcheon and Yosef Yarom. Resonance, oscillation and the intrinsic frequency preferences of neurons. Trends in neurosciences, 23(5):216–222, 2000.

[18] Nir Grossman, David Bono, and Edward Boyden. Methods and apparatus for stimulation of biological tissue, August 3 2017. US Patent App. 15/512, 556.

[19] Alan V Oppenheim, Alan S. Willsky, and S. Hamid Nawab. Signals and Systems (2Nd Ed.). Prentice-Hall, Inc., Upper Saddle River, NJ, USA, 1996.

[20] Frank Rattay. Analysis of models for external stimulation of axons. IEEE transactions on biomedical engineering, (10):974–977, 1986.

[21] E. M. Izhikevich and R. FitzHugh. FitzHugh-Nagumo model. Scholarpedia, 1(9):1349, 2006. revision #123664.

