## Supplementary information for "The Mechanics of Temporal Interference Stimulation" (from another version of this preprint: version 1, 2020-04-25)

### The Mechanics of Temporal Interference Stimulation: Supp info

This document contains supplementary information for “The Mechanics of Temporal Interference Stimulation” by Cao et al. Figures are large, so they appear on several pages.

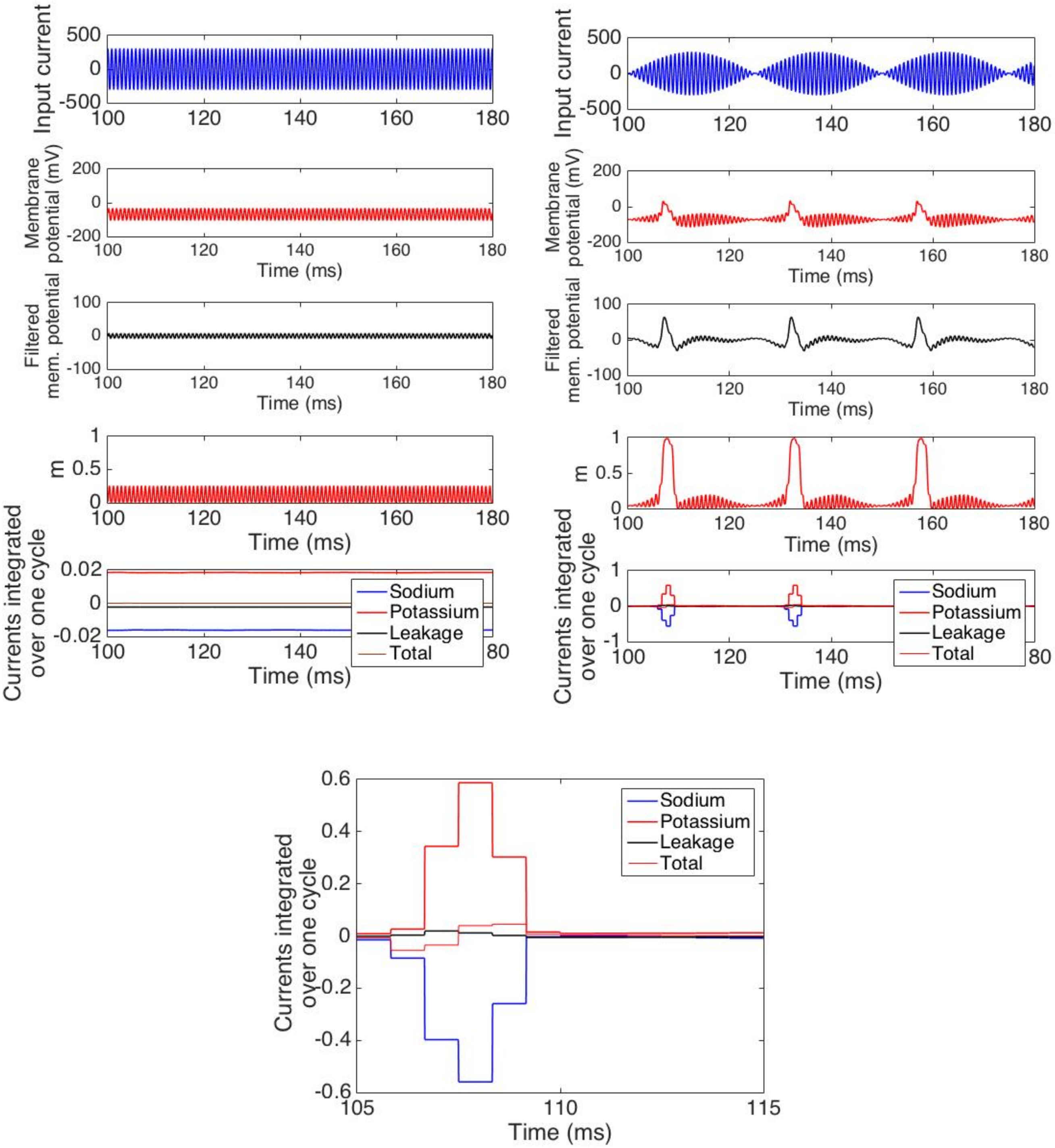


Fig. 1. If sodium channels are slowed down, TI stimulation is exhibited at lower frequencies: stimulation is observed with envelope modulated waveforms for 1200 Hz and 1240 Hz sinuosidal inputs, whereas a pure 1200 Hz sine wave (with twice the amplitude) does not cause stimulation. Current imbalance is shown in the lower figure. Maximum current amplitude is kept constant through Figures 1, 2, and 3.

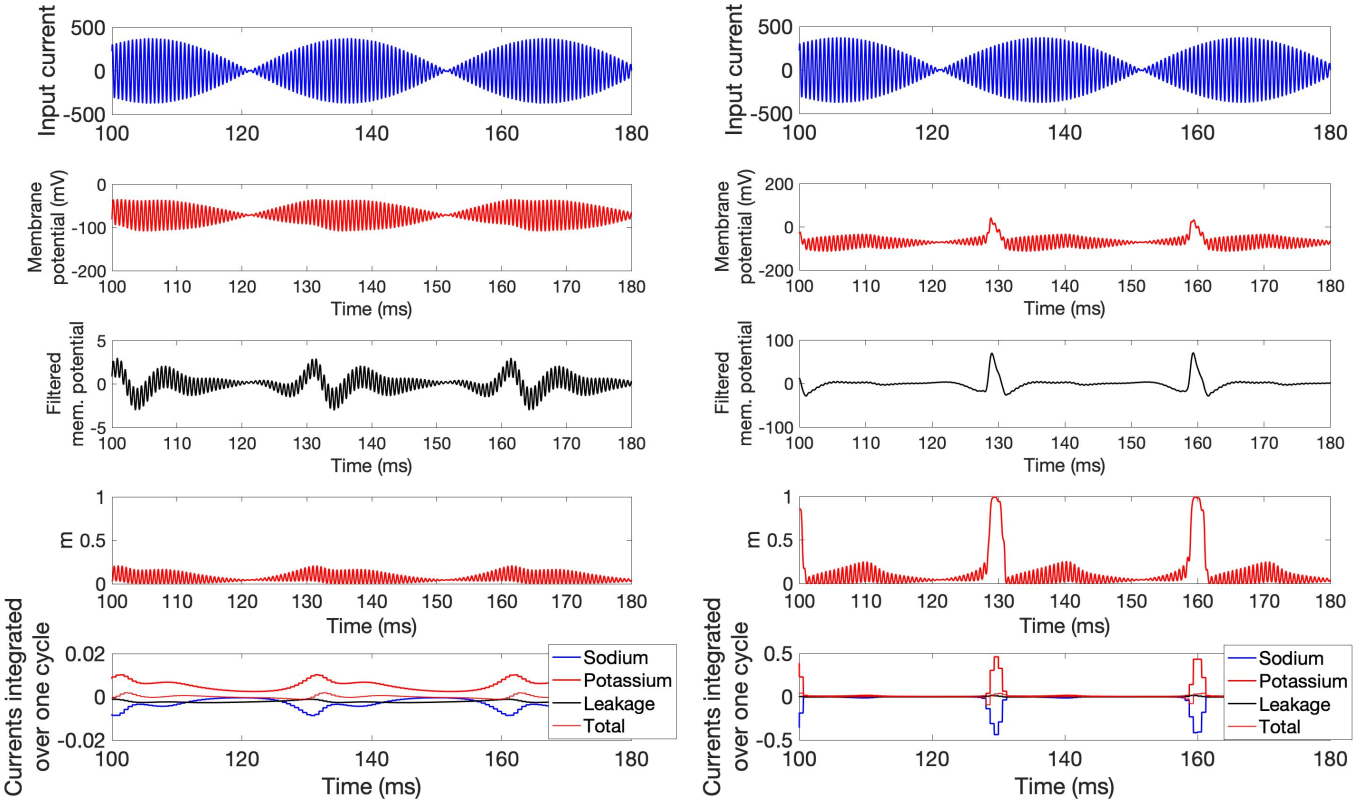


Fig. 2. Adding two sinusoids of frequencies 1600 and 1633 Hz, each of amplitude 187.27, a slowed down neuron (shown on left) does not fire, but the original Hodgkin-Huxley (shown on right) neuron does. Note the reduced amplitude of filtered membrane potential and the small $m$ values attained in the left figure (slowed down neuron) relative to that on the right.

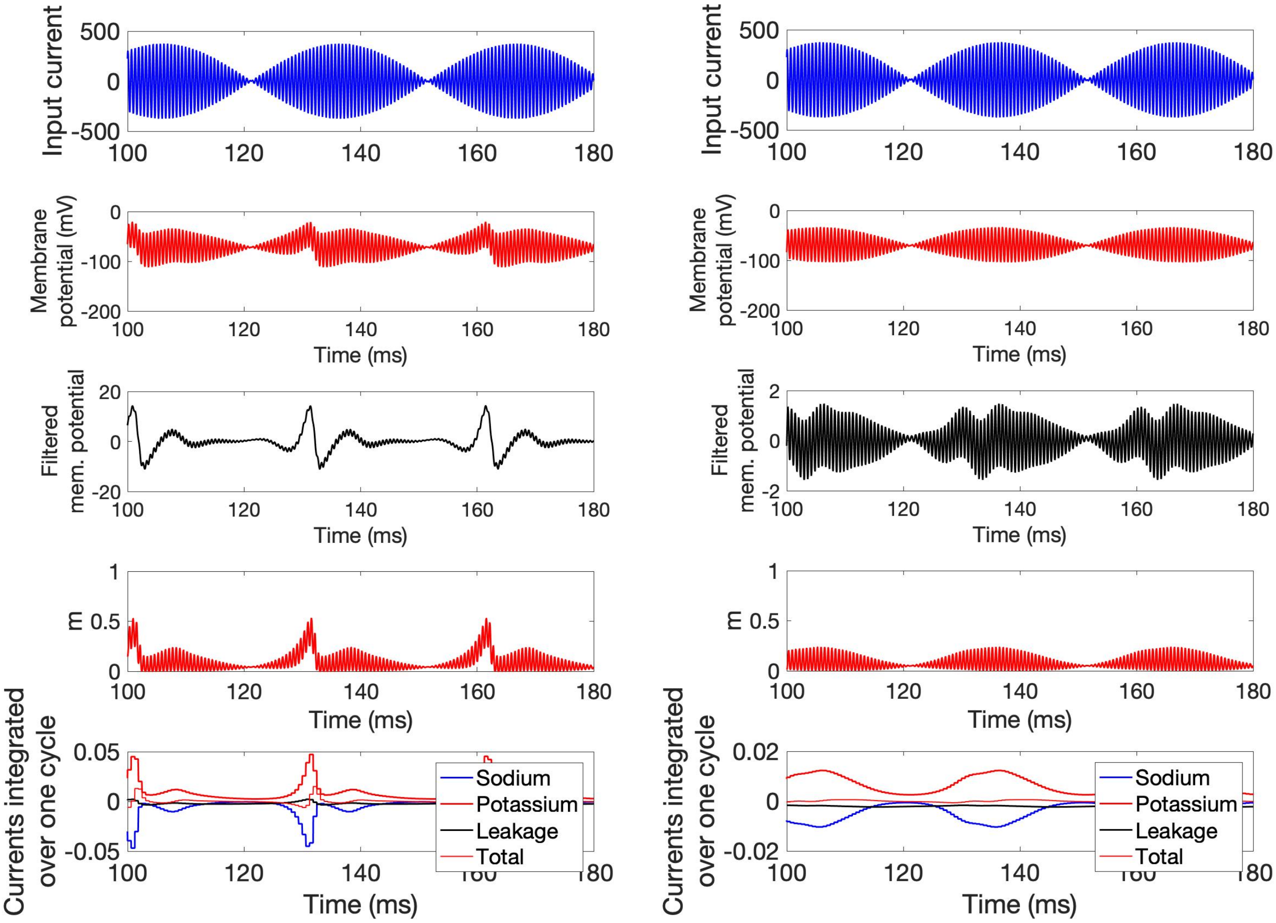


Fig. 3. Speeding up sodium inactivation parameter, $h$ (left), or Potassisum activation parameter, $n$, by 3x leads to the neuron not exhibiting TI stimulation at amplitudes and frequencies in Fig. 2.
